# Mapping the combinatorial coding between olfactory receptors and perception with deep learning

**DOI:** 10.1101/2024.09.16.613334

**Authors:** Seyone Chithrananda, Judith Amores, Kevin K. Yang

**Author notes:** Work done principally during an internship at Microsoft Research New England (Seyone Chithrananda). These authors contributed equally to this work (Judith Amores, Kevin K. Yang).

## Abstract

The sense of smell remains poorly understood, especially in contrast to visual and auditory coding. At the core of our sense of smell is the olfactory information flow, in which odorant molecules activate a subset of our olfactory receptors and combinations of unique receptor activations code for unique odors. Understanding this relationship is crucial for unraveling the mysteries of human olfaction and its potential therapeutic applications. Despite this, predicting molecule-OR interactions remains incredibly difficult. Here, we develop a novel, biologically-inspired approach denoted *MolOR* that first maps odorant molecules to their respective olfactory receptor (OR) activation profiles and subsequently predicts their odor percepts. Despite a lack of overlap between molecules with OR activation data and percept annotations, our joint model improves percept prediction by leveraging the OR activation profile of each odorant as auxiliary features in predicting its percepts. We extend this cross receptor-percept approach, showing that sets of molecules with very different structures but similar percepts, a common challenge for chemosensory prediction, have similar predicted OR activation profiles. Lastly, we further probe the odorant-OR model’s predictive ability, showing it can distinguish binding patterns across unique OR families, as well as between protein-coding genes or frequently occuring pseudogenes in the human olfactory subgenome. This work may aid in the potential discovery of novel odorant ligands targeting functions of orphan ORs, and in further characterizing the relationship between chemical structures and percepts. In doing so, we hope to advance our understanding of olfactory perception and the design of new odorants with desired perceptual qualities.

---

The sense of smell remains poorly understood, especially in contrast to visual and auditory coding. In vision, RGB maps directly to color, while in hearing, frequency maps to pitch. As a result, well-established maps coding for vision and hearing have been developed, such as the CIE color mapping (*1*) and Fourier space (*2*). In visual coding, well-established digitized maps allow any color to be represented by combining primary colors. Given the formation of a biological basis for vision through cone cells, the RGB mapping enabled the digitization of vision. In auditory coding, the single-axis frequency mapping enabled the ability to record and interpret sounds and to understand how different nerve endings in our ears are sensitive to different frequencies (*3*).

However, while we know that scent perception is mediated by binding to olfactory receptors (**Fig. 1A**), no such low-dimensional map exists for olfaction due to discontinuities in the mapping between the physical structure of an odorant molecule to its perceptual characteristics. While pitch increases with frequency, there exists no direct, well-characterized relationship between a molecule’s structure and its odor percept. This is often demonstrated using *Sell’s triplets* (*4*), which consist of three odorant molecules in which the pair with high structural similarity have low perceptual similarity while the pair with low structural similarity is perceptually similar (**Fig. 1B**). As a result, the standard chemoinformatic representations, such as molecular functional group counts, physical properties, or molecular fingerprints, are inadequate to map odor space.

**Figure 1:**
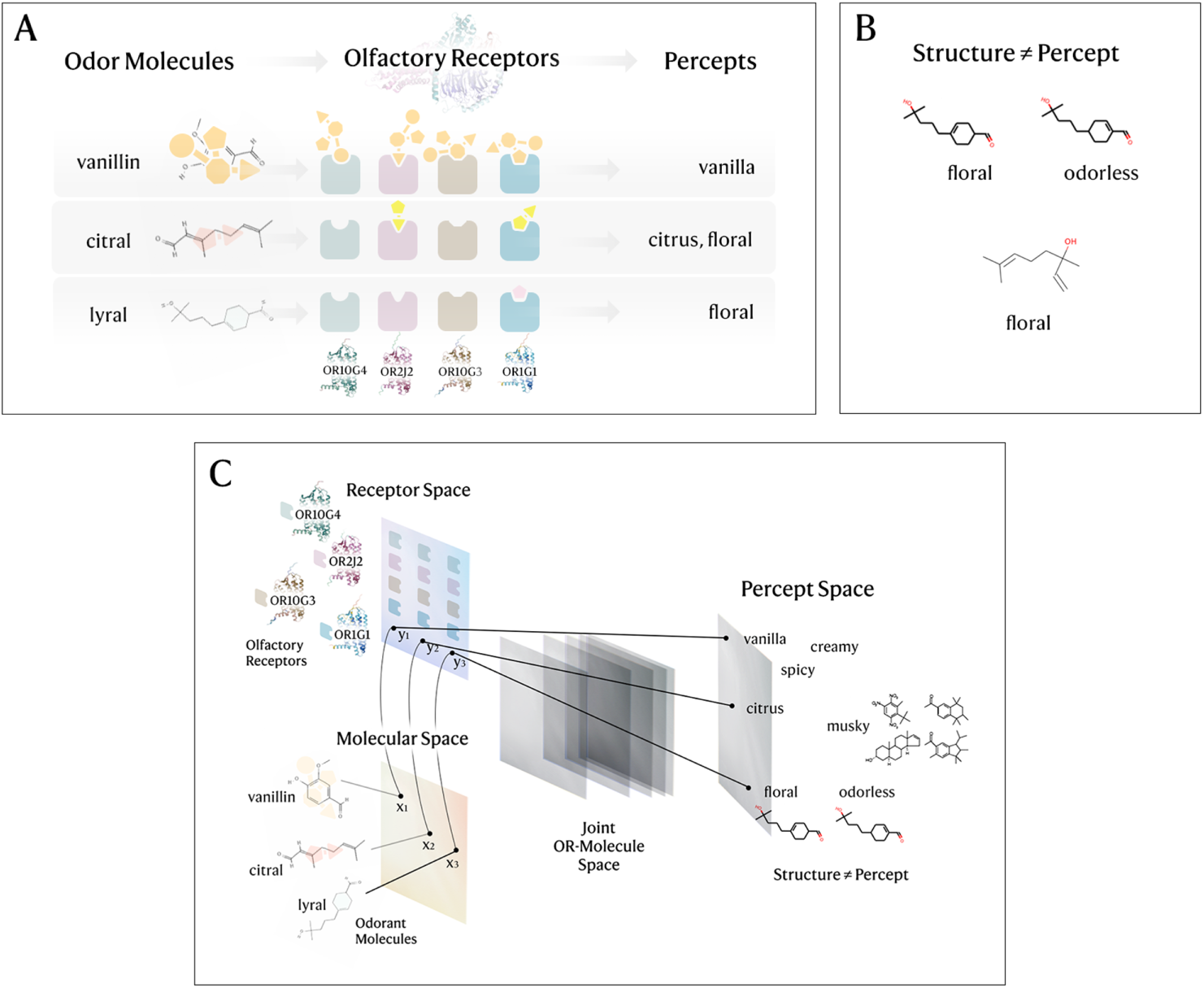
The molecular basis of olfaction. a) In the combinatorial coding hypothesis, odor molecules bind to one or more odor receptors, eliciting odor percepts. b) Sell’s’s triplets, in which molecular structure does not directly map to percepts based purely on physicochemical features. c) Inspired by the non-linearity of the molecular structure to odor perception relationship and the role of OR coding, we build hybrid models that learn a joint receptor-odorant space for percept prediction. Here, the “Joint OR-Molecule space” represents intermediate layers of the percept prediction model mapping the latent representation from the receptor predictions and molecular structure to the percept prediction.

Nevertheless, we now know the existence of the fundamental mechanisms underlying olfaction. Olfactory receptors (ORs) are G-protein-coupled receptors that line the hair-like cilia which protrude along olfactory sensory neurons’ (OSNs) dendrites. Unique to olfaction, each OSN expresses only one type of OR, and the combination of ORs (and therefore OSNs) activated produces a unique odor (*5*). The binding affinity between each of the 400 human olfactory receptors and an odorant molecule depends on features such as the shape and key functional groups of the odorant and the receptor’s binding site. Once an odorant binds to a receptor, receptor-mediated processing of mixture information occurs in the olfactory epithelium (*6*). The olfactory receptor undergoes a conformational change, activating a G-protein inside the cell. This triggers downstream signaling pathways that produce an electric signal that is then transmitted to the olfactory bulb in the brain, where further processing takes place. Under the combinatorial coding hypothesis (**Fig. 1A**), each combination of unique receptor activations codes for a unique odor percept (*7*), although other factors, such as the concentration of the odorant or temporal patterns may also affect how the brain deciphers these odorant responses. For example, the smell of freshly baked bread comes from over 500 volatile compounds, yet how these create its familiar scent remains unclear. Recent research (*8*) suggests that higher brain regions may use pattern recognition rather than an additive combinatorial code to form a perception.

Recently, researchers have applied deep learning to predict OR activation (*9*) and perceptual odor (*10*) from odorant molecular structure. On the other hand, prior work has utilized human OR activity and chemical features to predict perceptual data, measuring the ability of genetic variation in human ORs to affect odor perception (*11*). However, predicting OR activation and percepts separately does not leverage the mechanics of olfactory information flow.

Therefore, we develop a novel, biologically-inspired approach that first maps odorant molecules to their respective OR activation profiles and subsequently predicts their odor percepts (**Fig. 1C**). Our odorant-receptor activation model encodes residue-wise information from ORs with a pre-trained protein language model (*12*). Odorant atomic features are encoded by a graph neural network, and the OR and odorant encodings are combined to predict OR activation using an interpretable cross-attention mechanism. Despite a lack of overlap between molecules with OR activation data and percept annotations, our joint model improves percept prediction by leveraging the OR activation profile of each odorant as auxiliary features in predicting its percepts. We further note that sets of odorants within the same percept family yet diverse in molecular structure space have similar predicted OR activation profiles, which we investigate and visualize through OR-percept activation maps in Figure 5.

To further enable machine learning research in olfaction, we release our dataset of ∼6000 molecule-percept pairs processed from the pyrfume (*13*) package along with code and weights for our model. Our approach leverages biological mechanism to gain insight into the perceptual odor responses triggered by molecules, shedding light on the underlying relationship between molecular structure, receptor activations, and odor percepts.

## A dataset for molecule-OR binding-percept prediction

We set out to compile a dataset that captures the complex interactions between odorant molecules, olfactory receptors, and the resulting percepts. An ideal dataset for an end-to-end model of olfaction would contain both OR binding measurements and odor percepts for a set of diverse molecules, allowing end-to-end training and a deeper understanding of how molecular structures translate into unique olfactory perceptions. Because such a dataset does not exist, we explored the option of combining molecule-OR and molecule-percept datasets.

For molecule-OR data, we use M2OR (*14*), which combines experiments for 1237 unique receptor sequence variants for 11 mammalian species for a total of 46,700 molecule-OR pairs (**Fig. 2A**). We compiled a dataset of molecule-percept mappings by combining the Goodscents (*15*) and Leffingwell fragrance datasets (*16*), in a similar fashion to (*10*). These databases consist of expert ratings of percepts for individual odorants, with each molecule being linked to 1-15 percepts. One limitation of these percept datasets is the number of molecules per percept, as the joint dataset consists of only 669 unique, deduplicated percepts elicited by 5862 molecules. A majority of these percepts are elicited by fewer than 30 molecules while a few are elicited by up to 2000 molecules, creating an incredibly imbalanced regime (**Fig. 2B**). We further observe that a few select percepts (‘sweet’, ‘fruity’, ‘vanilla’) dominate majority of the labelled data, outlined by a handful of percepts with *>* 500 molecules (**Fig. S1**). Additionally, noise and various artifacts (such as the presence of odorous contaminants) when relying on human-rated odor perception to obtain ground-truth percepts is another major issue to overcome. Following (*10*), we removed percepts with fewer than 30 unique molecules in the joint Goodscents-Leffingwell (GS-LF) dataset, resulting in a multi-task binary classification dataset of 5862 molecules and 152 unique percepts. To determine the feasibility of training an end-to-end model, we computed the Tanimoto similarity coefficient (*17*) between molecules in the molecule-OR and molecule-percept datasets (**Fig. S3**). We observe that there are scarce pairwise structural matches between the datasets, with only 426 exact pairwise matches when mapping to canonical SMILES strings in both datasets. This makes training an end-to-end model jointly across both tasks difficult, so we chose to model both tasks separately.

**Figure 2:**
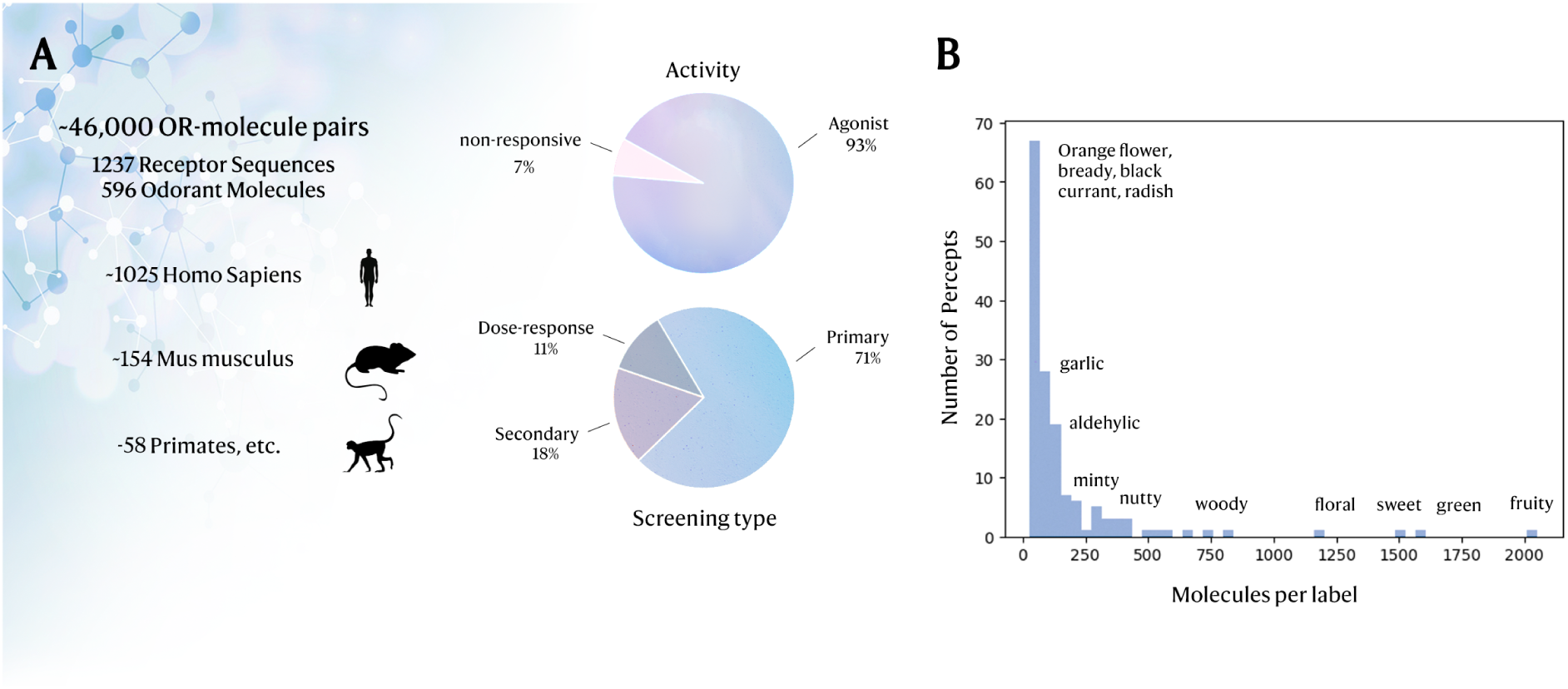
Assembling datasets for receptor-odorant interactions and odor perception. **A)** We utilize the M2OR (*14*) dataset, a diverse set of mammalian receptor-odorant pairs from various sources of experimental data (with varying degrees of noise). **B)** We highlight the imbalanced nature of both molecule-OR interactions and perceptual data, where only ahandful of percepts dominate the number of active odorant samples.

## A biologically-inspired machine-learning model of olfaction

Given the infeasibility of training a model to simultaneously predict OR binding and scent percepts for each molecule using existing datasets, we first train a model to predict molecule-OR binding using the M2OR dataset and then used predicted OR binding as an additional input when training a molecule-percept model using the joint GS-LF dataset. Full modeling details are provided in the Methods.

## Molecule-OR binding prediction

To design the molecule-OR model, we once again examine biology to glean desired inductive biases to overcome the low-data, noisy regime of molecule-OR binding. Molecule-OR binding is mediated by chemical interactions between functional groups in the odorant and specific amino acids in the receptor. Experimental structures of molecule-OR complexes show that often only a few residues in the OR and a few atoms in the molecule interact directly. We therefore develop a new model architecture (**Fig. 3A**) that uses a cross-attention mechanism to project atom-wise and residue-wise embeddings into a single, joint embedding space, allowing the model to learn to attend to the most important residues and atoms without needing to explicitly identify the binding site (due to the lack of structural data for ORs).

**Figure 3:**
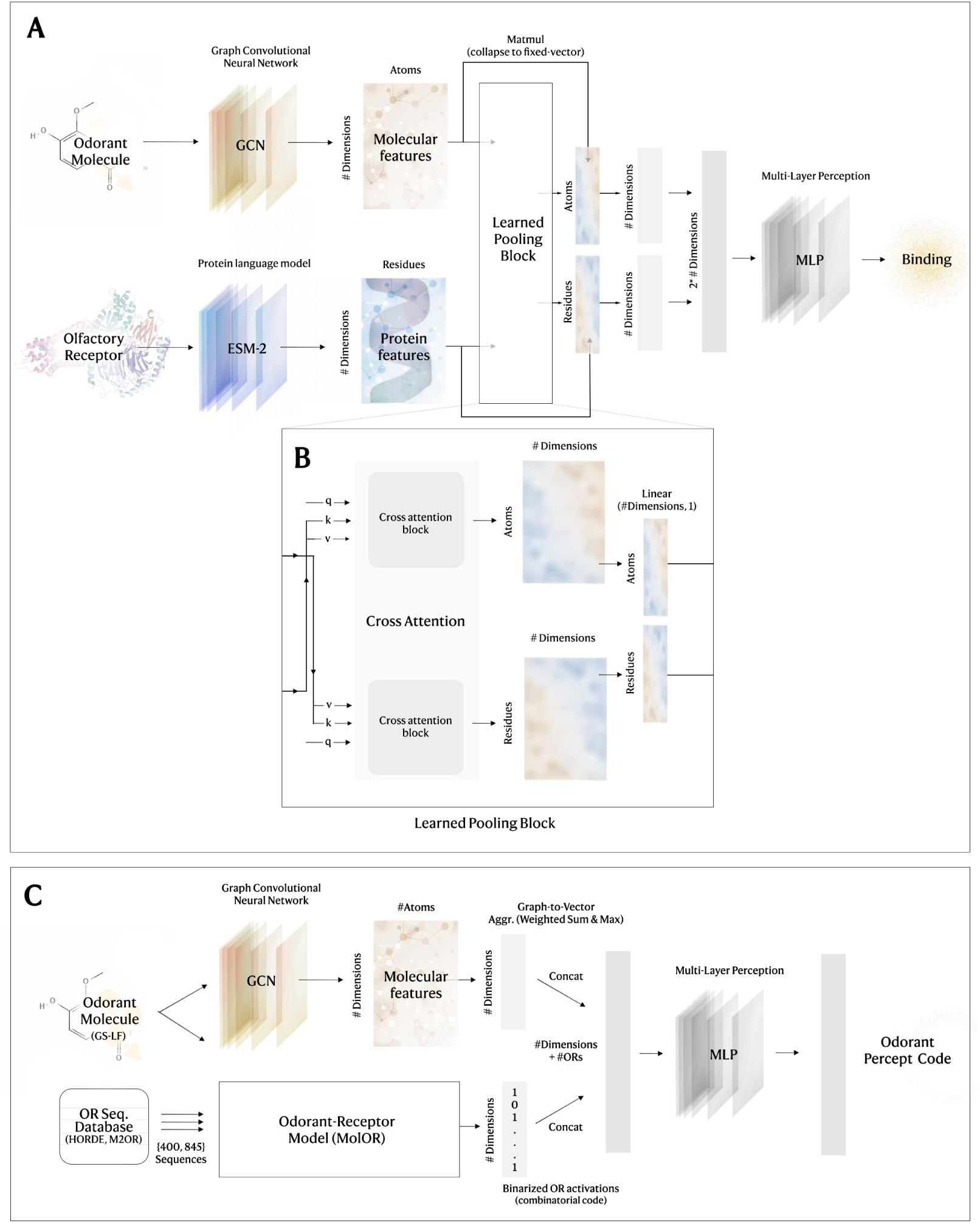
**A**) Receptor-odorant model (MolOR) integrates protein language models with geometric learning to fuse learned information from both molecular and protein representations. **B) Learned pooling block** in which cross-attention blocks produce per-atom and per-residue vectors, which act as *weights* to aggregate the 2D molecule/protein tensors produced by the backbone models (GCN, ESM-2). **C) Odor perception model** We use MolOR to generate *syn-thetic* activations for olfactory receptors in HORDE, and use these in tandem with the molecular features from a GCN to predict odor perception across a list of 152 percepts from fragrance databases.

We use a graph convolutional network (GCN) (*18*) to learn the atom-wise molecule embeddings; GCNs leverage the fact that molecules consist of atoms connected by bonds. In parallel, we use residue-wise OR embeddings from the pretrained ESM-2 protein language model (PLM) (*12*). PLMs are pretrained on tens of millions of protein sequences to capture evolutionary and structural information, and so the per-residue embeddings have been shown to act as a meaningful prior for protein structure and property prediction models (*19*). The learned pooling block then produces a set of interpretable cross-attention scores that determine how much to weight the individual residues and atoms in both embeddings, which are then applied to the molecule and protein embeddings to obtain the final protein and molecule embeddings (**Fig. 3B**). **Table 1** shows that learned pooling with cross-attention between GCR and ESM-2 embeddings outperforms the GCN by itself, a simple concatenation of the atom-wise GCN and residue-wise ESM-2 embeddings, and using a randomly-initialized version of ESM-2, showing that both protein sequence pretraining and the learned pooling improve performance. We also benchmark against leading models for the drug-target interaction problem, such as PerceiverCPI (*20*), CPI architectures used in enzyme-substrate interaction prediction (*21*), as well as odorant-receptor model OdoriFy (*22*). PerceiverCPI utilizes a series of Perceiver IO-like cross-attention steps on a) multiple embedding architectures (MPNN, Random Forest) across different molecular representations (graphs, fingerprints), and b) final molecular and protein per-residue embeddings (encoded using a series of 1D CNN layers) (*23*). The enzyme-substrate interaction mdoel detailed in (*21*) combines pretrained neural networks which extract features from the substrate and enzyme to be fed into a top-level feed forward model for activity prediction, akin to the GCN w/ PLM embeddings. They concatenate Morgan fingerprints with mean-pooled ESM-1b protein language model embeddings through a feed-forward network, representing a feature-concatenation approach to compound-protein interaction prediction without learned cross-attention. OdoriFy uses two BiLSTMs with linear transformation heads, which takes as input one-hot encodings of the OR sequence and odorant SMILES, respectively. The modality-specific embeddings are concatenated before being passed through a classification head (*22*).

**Table 1:** Odorant-OR binding prediction. Uncertainties are standard deviations across 5 random seeds. **Model** denotes architecture, including encoder type and whether receptor-level features were used. **Random/Pre-trained** denotes what type of protein sequence model was used, whether it was pre-trained on a broader corpora, and its model size. **Pooling** ablates mean-averaged embeddings compared to a learned residue-level aggregation. **Loss Type** denotes the sample weighing scheme used to train the model. We evaluate a multi-task GNN where binding to each unique OR is posited as a “task”, a GCN with mean-aggregated pLM embeddings, three existing models (Odorify (*22*), PerceiverCPI (*20*), Enzyme-substrate FFN (*21*)) and our architecture with a different encoder architecture (MPNN), against different ablations of MolOR.

| Model | Random/Pre-trained | Pooling | Loss Type | Test AUROC |
| --- | --- | --- | --- | --- |
| Multi-task GNN | N/A | N/A | Unweighted | $65.88 \pm 0.27$ |
| GCN w/ PLM emb. | Pre-trained (ESM-650M) | Mean | Unweighted | $78.92 \pm 0.37$ |
| MolOR | Random (ESM-650M) | Learned | Unweighted | $85.49 \pm 0.93$ |
| MolOR | Pre-trained (ESM-650M) | Learned | Weighted | $86.78 \pm 0.80$ |
| **MolOR** | Pre-trained (ESM-650M) | Learned | Unweighted | **$88.97 \pm 0.30$** |
| MolOR | Pre-trained (ESM-3B) | Learned | Unweighted | $88.10 \pm 0.38$ |
| MolOR (MPNN enc.) | Pre-trained (ESM-650) | Learned | Unweighted | $86.20 \pm 0.04$ |
| PerceiverCPI (20) | Random (CNN) | Learned | Unweighted | $85.47 \pm 0.42$ |
| Goldman et. al (FFN) (21) | Pre-trained (ESM-1b) | Mean | Uneighted | $86.82 \pm 0.53$ |
| OdoriFy (22) | Random (BiLSTM) | Learned | Weighted | $77.73 \pm 0.46$ |

We train molecule-OR binding models using binary cross entropy (BCE) losses. However, motivated by the presence of several sources of biases in the data, we investigated the performance of the weighted loss scheme proposed by (*9*), which accounts for label noise coming from high-throughput screening, class imbalance, and the combinatorial nature of the task. We find that while the weighted loss performs considerably worse at matching receptors to molecules, it slightly outperforms the unweighted BCE loss when the molecule-OR model is used to generate synthetic OR activations as context for percept prediction. We speculate that the weighted loss places more emphasis on underrepresented binding events via the ligand/receptor data frequency terms in the pairwise sample weighing scheme, which slightly reduces raw binding accuracy but produces a binding model whose outputs are more generalizable for downstream percept prediction. We fully describe the weighted loss in the Methods.

Lastly, we construct evaluations to measure reliance on train-test overlap, and potential leakage between training and test sets. We perform leave-one-subfamily-out (LOSO) experiments, in which all member sequences of an OR subfamily (OR2J) are withheld during training of MolOR models. We then evaluate MolOR’s ability to predict responses for the held-out OR2J subfamily receptors. Evaluating OR2J sequences is a distinct challenge for two reasons; the class ratio of OR2J receptor-odorant pairs in M2OR is three times more biased in favor of agonists compared to the broader dataset, and has a two-fold higher ratio of pairs coming from dose-response screens (*EC*_50_). Despite this, results show that ROC-AUC on withheld functional receptors across seeds remains relatively high (75.05 ± 0.52), suggesting that the model leverages sequence features from the pretrained model embedding rather than just memorizing the distribution of labels related to a specific receptor sequence, screening type, or odorant molecule in the train set.

## Molecule-percept prediction

Inspired by the theory underpinning olfactory information flow, we concatenate a panel of predicted binary molecule-OR binding states to odorant molecular embeddings from a graph convolutional model to predict the elicited percepts. This embedding forms a joint OR-molecule space learned by the MLP percept prediction head. Including predicted binding states consistently improves percept predictions (**Fig. 4**) over using the graph convolutional network on odorant structures alone, regardless of the OR database used (Table S1). To evaluate this, we test using predicted MolOR activations against two olfactory receptor databases, M2OR, and HORDE (*24*). HORDE mined and elucidated the nearly complete human olfactory subgenome, searching the genome draft with gene discovery algorithms to identify 900 olfactory receptor genes and pseudogenes. This makes it a rich dataset for characterizing if the model can differentiate binding patterns across human OR gene families. Thus, we speculated whether it would provide a more useful prior for distinguishing odor perception.

**Figure 4:**
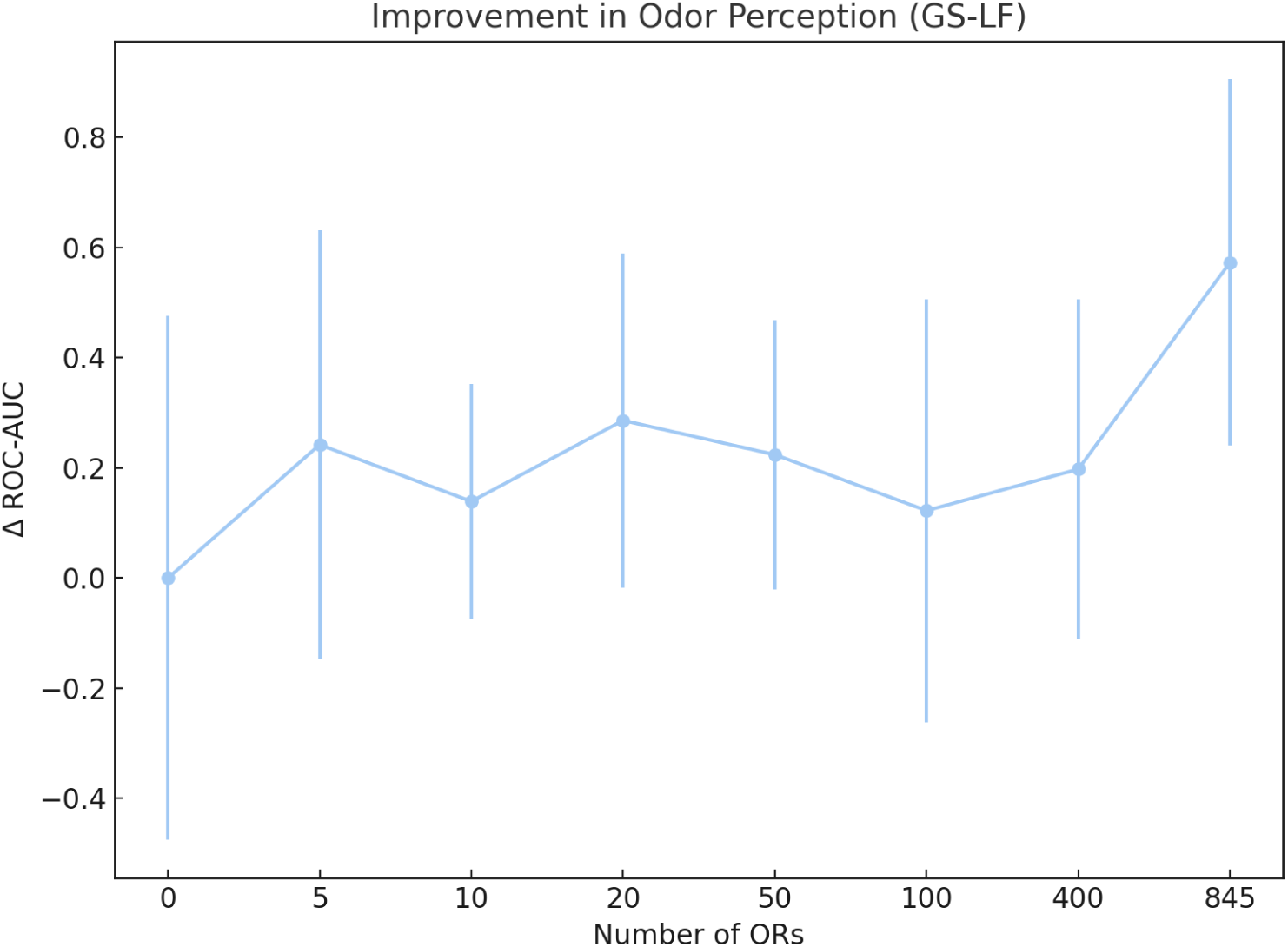
Improvement in odor percept prediction when using predicted MolOR activations for ORs in HORDE. We measure mean AUROC over 5 seeds across each ablation.

**Figure 5:**
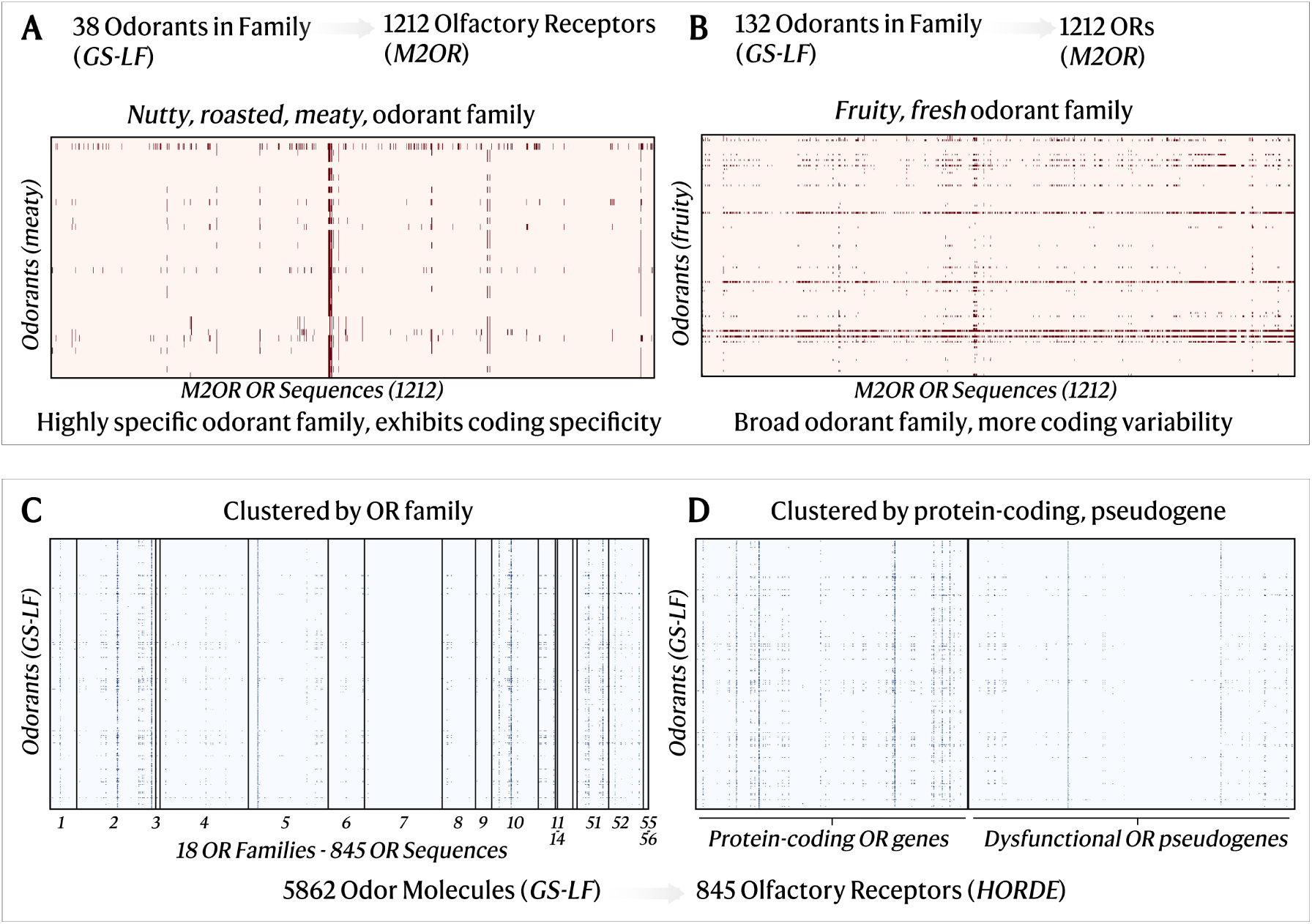
MolOR predictions for ORs in M2OR, HORDE. **A)** Nutty, roasted, meaty odorants. **B)** Fruity, fresh odorants. The ORs in **A-B** are sorted in descending order from left to right by how frequently they are repored as active in the M2OR dataset. **C)** Activations across odorants with human ORs grouped by family annotations in HORDE. **D)** Activations across odorants for human OR genes vs pseudogenes in HORDE.

We empirically find that percept prediction improvement against the baseline peaks using the weighted loss in the binding model with either HORDE or M2OR. Using all 845 OR activations in *HORDE* or a panel of the top 400 most-commonly bound OR sequences in *M2OR* (**Table S1**), we observe a statistically significant tren of improvements to percept prediction as we increase the number of OR predictions (Jonckheere-Terpstra *p* = 0.010, **Fig. 4**, **Table S1**), though the absolute effect size is modest. We also benchmark against the MPNN architecture defined in (*10*), as well as the performance difference between using synthetic predictions from weighted and unweighted MolOR models (**Table 2**).

**Table 2:** Goodscents-Leffingwell percept prediction, by model type and synthetic data usess.

| Model | MolOR Loss Type | OR Database | Test AUROC |
| --- | --- | --- | --- |
| MPNN - baseline (10) | N/A | N/A | $86.32 \pm 0.54$ |
| GCN (Pseudogenes only) | Weighted | HORDE | $87.04 \pm 0.34$ |
| GCN (0 ORs) | N/A | N/A | $87.04 \pm 0.48$ |
| GCN (400 ORs) | Unweighted | M2OR | $87.25 \pm 0.22$ |
| **GCN (400 ORs)** | **Weighted** | **M2OR** | **$87.57 \pm 0.49$** |
| GCN (845 ORs) | Unweighted | HORDE | $87.20 \pm 0.42$ |
| **GCN (845 ORs)** | **Weighted** | **HORDE** | **$87.62 \pm 0.33$** |

We also investigate whether pseudogenes, ORs which are similar in sequence space to functional ORs but crucially **nonfunctional proteins** are contributing to percept prediction capabilities. Here, we aimed to see whether more functionally irrelevant interactions were artificially boosting percept prediction performance, due to their proximity to functional human ORs. We find that the performance using only pseudogene predictions on GS-LF (87.04 ± 0.34) is nearly identical to the baseline of using no OR logits (just the base GCN model) for percept prediction. In comparison, using predictions against the entire HORDE database of receptors *(386 intact ORs, 459 psuedogenes)* performs 0.6% better (87.62 ± 0.33). This suggests that the model essentially discards the pseudogene receptor-odorant predictions (which are mostly predicted as inactive, as seen in 5).

Lastly, we measure the contribution of progressively increasing the number of receptors from which we pool predicted binding states as features for percept prediction, as detailed in **SI Table S1.** Additionally, we run a paired t-test against the baseline percept model (no OR predictions) and model leveraging activation’s across all of *HORDE* (845 ORs). We compare ROC-AUC performance across 5 seeds for both, and produce a t-statistic of 5.91 with p-value 0.004, making the mean difference between the two ablations significantly different from zero. To test whether the overall scaling trend is robust, we additionally applied a Jonckheere-Terpstra test for monotonic ordering across all 8 ablation levels (0 to 845 ORs), yielding *p* = 0.010.

## Biologically-inspired models uncover potential combinatorial codes for odor families

Inspired by the combinatorial coding theory of olfaction, we investigate whether our model can elucidate unknown combinatorial codes linking olfactory receptors to specific odor percept families. One major caveat to the universal combinatorial coding theory in olfaction is that different cultures often perceive smell in strikingly different ways due to differences in experience (*25*), which has many implications for the viability of a model trained on fragrance databases with an unknown distribution of cultural backgrounds across the reviewer base. However, recent work has found that while individuals vary as to which odors they find pleasant or unpleasant, their culture only explains 6% of the variance (*26*): pleasantness may be shared across cultures. Therefore, we investigate if our model consistently predicts reoccurring activated ORs for molecules that are both pleasant or unpleasant but otherwise diverge perceptually (eg: ‘vanilla’, ‘creamy’). We also ensure that we account for OR promiscuity, ensuring that ORs which simply activate a large proportion of the odorant space are not misidentified as part of the ‘unique combinatorial code’ for a given percept.

First, with the base MolOR model, we predicted OR activation’s from the molecule-OR model for 1237 olfactory receptor sequences seen in the M2OR database, across all 5862 odorants in GS-LF (S2). Out of 1237, 25 olfactory receptor sequences are predicted as responsive across over 30% of the Goodscents-Leffingwell database (GS-LF). To better probe specificity/selectivity across the combinatorial code of odor-families, we filter these 25 sequences out for generating OR-odor family plots. We observe that for 9/25 of these promiscuous OR receptor sequences, a BLAST (*27*) search with E-threshold 10 shows they have homology to the mammalian OR2J3 olfactory receptor family, which are highly enriched in the M2OR training data as high-quality, activated receptors from dose-response screens. We assume that this is due to the upweighting of pairs from high-quality data types. Additionally, we filtered out odorant molecules predicted as *broadly* active (i.e., any molecule in GS-LF for which *>*30% of ORs in M2OR are predicted as active). We observe that for specific subfamilies of percepts in fragrance wheels, our models predict specificity in distinct receptor activations (ex: ‘meaty’, ‘warm’), but for broader classes (‘fruity’) containing many odorants, we see more variance in predictions, with odorants binding to many different classes of receptors (**Fig. 5A-B**). To assess statistical sensitivity of synthetic percept-receptor associations, we also quantify the specificity of these associations against a null model distribution (via random permutations of the percept labels across molecules). We provide details on these experiments in SI.

A major limitation of such interpretability analysis identifying OR-percept associations in a model-guided fashion, is that enriched ORs may not match reality without experimentation. To probe whether this model can identify percept family-associated ORs, we examine the proportion of molecules part of the “nutty” percept family (“nutty, roasted, meaty”) that are identified by the top OR enriched across that family. This can also be seen as computing the precision of using the top “nutty” family-enriched OR to identify molecules that are likely to have the percept. Using the weighted-MolOR models’ predicted activations (as they are the same features used in our percept prediction model), we apply the same filter for dropping promiscuous ORs as done presently in **Fig. 5A**, where any OR predicted to activate more than 30% of GS-LF is dropped. We apply a similar for odorants (any odorant from GS-LF that binds to more than 20% of ORs in M2OR is dropped), yielding a filtered OR predictions matrix of size (4355 molecules, 1105 ORs). Despite using synthetic MolOR predictions on unseen molecules, we find OR5K2 exhibits the highest precision-recall curve for both the broader “nutty” percept and the associated specific family. Across all nutty molecules (*n* = 261), ORK52 has a precision of 36.4% (90/247 molecules predicted to bind with probability *>* 50%). Across the specific family, OR5K2 is predicted to activate 74.1% of all molecules, while only 5.2% of non-family-specific molecules bind to the receptor.

Additionally, using the HORDE (*24*) database, we examine MolOR-predicted OR activation maps against all odorants in the previously constructed Goodscents-Leffingwell dataset. First, we examine activity by OR family across non-promiscuous odorants in GS-LF (molecules that are predicted as activating *<*30% of HORDE receptors). Across over a dozen unique OR families, we see unique binding patterns, with certain families (ex: OR family 7) exhibiting nearly no binding activity (containing primarily “orphan” ORs), while others exhibit far more broadly-tuned activity (**Fig. 5C**).

Next, using binary predicted activations from the unweighted MolOR model, we plot variances in the frequency of activation’s for functional OR genes as compared to pseudogenes (**Fig. 5D**). We aim to see if the model is able to discern that non-functional ORs, despite having homology to protein-coding genes, cannot be “activated”. This task makes for an interesting challenge as more than half of the HORDE mined olfactory receptors are pseudogenes, which has been theorized to be due to a low selection pressure to maintain ORs that are functionally redundant or detect odours no longer relevant for a species’ fitness (*28*). Additionally, only some pseudogene fragments are translated, indicating odorant binding and activation should be much less frequent (*29*). Examining the predicted activation’s per odorant, we see a visible shift in the distribution of activated receptors per odorant for functional OR genes as compared to pseudogenes. The median number of activated protein-coding ORs is 3 for each GS-LF odorant, and only 1 across pseudogenes (Mann-Whitney U-test, 35.62, p *<* 0.01).

## Discussion

The field of olfaction has made enormous strides in understanding, through advances in biochemistry, psychology, and neuroscience, how diverse stimuli induce the perception of up to one trillion unique odors (*30*) (*31*). However, the olfactory system, responsible for our sense of smell, poses a unique challenge. Unlike the well-established mappings between wavelength and color or frequency and pitch, the relationship between chemical structures of molecules and their olfactory percepts, or odors, remains poorly understood. Thus, there is no concrete mapping from molecules to their scent. However, previous work has shown that receptor mechanisms, or the combination of olfactory receptors activate by odorants, plays a key role in governing perception. This combinatorial code, representing the responses of receptors elicited by olfactory stimuli (odorant), is effectively an input code for the neural deciphering of odor sensing. Odorant molecules form the *molecular space*, which are then recognized by an OR repertoire and mapped to *receptor code space* (*32*). Information from this receptor code space is what is processed by the brain and mapped to *odor percepts*.

Graph neural networks (GNNs) have exhibited promise in learning robust representations of molecules, especially odorants, leading to state-of-the-art performance in predicting the percepts of molecules (*10*). Driven by the biological information flow underlying odor perception, advancements in geometric learning for odorants and protein modeling of olfactory receptors, we aimed to combine the previously disjoint views of olfaction: modeling receptor-odorant interactions, and modeling the molecule-percept map. Concretely, we found that co-embedding GNN odorant molecule features with predicted OR activations results in more accurate percept prediction than using odorant molecule features alone. Beyond predicting OR-binding, our approach also offers insights into the underlying relationship between molecular structure, receptor activations, and odor.

There exists several limitations and opportunities for follow-up work. The concentration of individual odor molecules has a significant impact on the perceptual profile it produces, but our current model does not account for this, and so can only predict odor detection beyond some threshold, but not *intensity*. Additionally, our model does not account for the odor profiles of mixtures of compounds, which can elicit novel percepts. One other direction for future work is accounting for odorous contaminants, especially those that have not been developed for use in fragrance applications. Future work could incorporate mixtures by accounting for non-bonded interactions and chirality of individual molecules, as done in (*9*).

Lastly, we believe one of the most-needed directions for furthering progress in machine learning in olfaction is collecting well-curated, diverse datasets for interactions between odorant molecules, receptors and odor percepts. Currently, our biggest limitation lies in the underlying OR–odorant binding data: assays span noisy single-concentration primary screens to higher-confidence dose–response (*EC*_5_0) measurements collected across heterogeneous experimental protocols and cell systems, and the receptor–odorant matrix is both highly imbalanced and sparsely sampled. In addition, there is almost no molecular overlap between the OR binding datasets and the perceptual datasets we use (Fig. 2B, Fig. S3), which prevents training a truly end-to-end model of combinatorial coding and odor perception. Future progress will therefore depend on larger, more standardized OR–odorant panels with dose–response measurements and better overlap with psychophysical datasets.

With the demonstrated potential of GNNs in predicting olfactory receptor binding and perceptual odor responses, we anticipate that machine learning can play a pivotal role in olfactory research, much like it has done for vision and hearing. Such work towards building model-driven maps of perceptual and receptor space may open up possibilities to enhance our understanding of olfactory processes, with potential applications in diverse fields such as human nutrition, fragrances, environmental impact assessment, diagnostics and sensory neuroscience.

## Acknowledgements

The authors thank members of Microsoft Research New England for helpful discussions over the course of this work. We thank Hattie Chung, Neil Malinar, Avish Vijayaraghavan, David Alvarez Melis, Sadhana Lolla, and Michelle Si for ideas and discussions that helped us improve our work. We also thank Machine Learning at Berkeley for compute resources for data processing and model training. S.C. was supported by the Masason Foundation, the Mercatus Center for the Emergent Ventures Fellowship, and New Science as part of the Computational Life Sciences microgrant.

## Declarations

A provisional patent has been filed on some of the technologies referenced to in this work by Microsoft Corp. K.K.Y. and J.A. are employees of Microsoft Corp., while this work was conducted by S.C. as part of an internship in 2023.

## Author contributions

Conceptualization: S.C., J.A. K.K.Y.; Methodology: S.C., J.A. K.K.Y.; Software Programming: S.C.; Validation: S.C., J.A. K.K.Y.; Formal analysis: S.C., J.A. K.K.Y.; Resources Provision: J.A., K.K.Y.; Data Curation: S.C., J.A. K.K.Y.; Visualization: S.C., J.A. K.K.Y.; Writing: S.C., J.A. K.K.Y.; Supervision: J.A., K.K.Y.

## Resource availability

Code, datasets, and models related to this study are publicly available at https://github.com/microsoft/olfaction.

## Methods

### Dataset assembly

For the molecule-OR dataset, we begin with 46,700 molecule-OR pairs from the M2OR dataset, which contains a subset of possible pairings between 596 molecules and 1237 unique receptor sequences. For the multiclass-GNN setting, we cluster the 1237 sequences (many of which are variants of a wildtype receptor) into a set of well-known olfactory receptor IDs by running BLAST on each sequence against UniProt and mapping it to the closest of‘canonical’ OR (yielding 571 OR “labels”). For all models that incorporate receptor sequence information, we simply use the 46,700 provided molecule-OR pairs in the original training set as individual, receptor-specific samples.

For the molecule-percept dataset, we begin with the Goodscents (GS) dataset as compiled by the pyrfume package (*13*), which maps molecules to a set of percept labels. Leffingwell (LF), on the other hand, uses freeform text. The set of unique odor descriptors across both datasets were canonicalized using the GoodScents ontology of descriptors and overlapping molecules inherited the union of odor descriptors from the two sources. After merging the two datasets, we infill missing percept labels against the set of 669 labels for odorants with NaNs. Percepts with fewer than 30 positive molecules were discarded, resulting in a set of 152 percepts.

### Embedding odorant molecules with graph neural networks

We use graph convolutional networks (*18*) to learn molecule embeddings. The network consists of several graph convolutional layers, which iteratively transform the node features by incorporating information from neighboring nodes. Specifically: define an adjacency matrix *A* representing the connections between nodes in a graph, matrix *X* representing the row-wise stacked node feature vectors, and matrix *D* representing a diagonal matrix where any diagonal element *D_i,i_* stores the total number of neighboring nodes to node *i* (1 if just itself). The layer will set diagonal elements to 1 such that during each node feature vector update, the node itself is passed in and maintained in the updated embedding. From there, the adjacency matrix is normalized with respect to the degree matrix, aggregated against the raw node feature vectors (the message passing step), before then updating the node feature vectors using weight matrix *W* with respect to the output dimension size of the GCN layer.

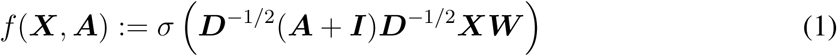

### Embedding odorant receptors with ESM-2

For incorporating protein information, we utilize the ESM-2 protein language model (model id: `esm2 t33 650M UR50D`), a 650M-parameter BERT-like model trained to correctly recover sequences with corrupted tokens. We hypothesized that because PLM representations encode structure, using the per-residue embeddings of OR sequences can inject a structural prior into the molecule-OR model. As a result, we generate per-residue embeddings for all 1237 novel receptor sequences in the M2OR dataset using ESM-2.

### Predicting percept or OR binding from molecule alone

The GCN produces atom-wise embeddings which must be aggregated into a fixed length vector in order to make a prediction about the entire molecule. For the molecule-only OR binding model (in the multi-task setting, using no implicit sequence information), after refining node representations using the GCN layers, we use a weighted sum and max aggregation to aggregate the per-atom representation. The fixed-length vector is fed into a set of dense layers, outputting a set of predictions for all OR labels. For the molecule-only percept prediction model, we similarly aggregate the atom-wise embeddings via a weighted sum with learnable weights and via max pooling over the set of nodes and then concatenate the output of the two aggregations, which is then fed into an MLP for final prediction.

#### Training details

For the canonical GNN baseline (for percept-prediction), we use two GCN layers, with a hidden size of 256, and a two-layer MLP with hidden dimension 128 for the prediction head. We employ the same GCN + prediction head on the multi-task M2OR setting. Both models use a learing rate of 2*e* − 2, on a batch size of 32 molecules, with residual connections between each GCN layer. We train for 100 epochs, with a patience of 30 epochs for early stopping, taking the best checkpoint before the patience is reached to benchmark on test. All models are trained on a single A100 GPU. We employ both random and stratified splits (80/10/10), but find that both sample an similar amount of samples from different data/screening qualities for M2OR, opting for random instead.

### Cross-attention model for molecule-OR binding prediction

We develop a learned cross-attention module to aggregate molecule and OR embeddings. The cross-attention block acceptd inputs from two different modalities; the query *Q* comes from modality 1 and key, value pairs *K, V* come from the other. *d_k_* is defined as the key’s embedding dimension:

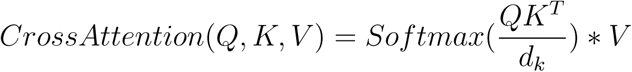

For both the protein embedding of shape (`# residues, protein embedding dimension`) and molecular embedding with shape (`# atoms, molecule embedding dimension`), we define a set of linear layer transforms for mapping to the input query, key, and value tensors for cross-attention, *f_Q_, f_K_, f_V_*. The linear layers map both molecule and protein embedding tensors to the same embedding dimension (for `esm2 t33 650M UR50D`, 1280). The two cross-attention blocks, for input tensors *p, m* are defined as:

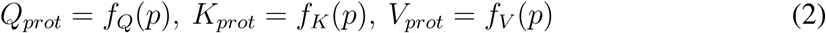

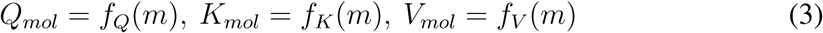

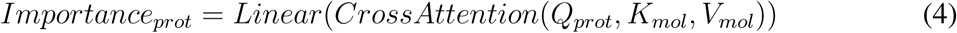

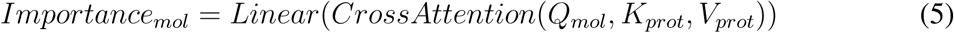

Where the outputs of the two cross-attention blocks are mapped to fixed vectors of size *L* using a linear layer, where *L* ∈ (#*residues,* #*atoms*), representing residue and atom importance scores. Using this cross-attention mechanism, the model generates an attention map that defines the set of scores from which the molecular and protein feature maps can use to do aggregation and obtain a vector embedding.

#### Training details

We employ the GCN + PLM cross-attention strategy (denoted *MolOR*) on the M2OR dataset of receptor-odorant pairs. We use the same GCN encoder and MLP prediction heads as the baseline above for fair comparison, with a dropout rate of 5%. Thus, the only components in the architecture that are changed are the integration of protein embeddings for receptor sequences, and the aggregation policy learned using cross-attention (or mean-averaged, as a baseline for protein embeddings). We similarly train for 100 epochs, with a patience of 30 epochs for early stopping, taking the best checkpoint before the patience is reached to benchmark on test. All models are trained on a single A100 GPU, and employ random splits (80/10/10), as we find that it samples a balanced amount of samples from different data/screening qualities for M2OR.

### Weighted Loss Scheme

For majority of the trained models, we utilize a vanilla binary cross entropy scheme for odorant-receptor binding prediction. However, we experiment with the weighted loss scheme proposed by (*9*), which accounts for variance in sample equality from the experiment type (**sample quality weights**), class imbalance due to most odorant-receptor pairs being non-responsive (**label imbalance**), and **pair-imbalance** accounting for up-weighing olfactory receptors that have been tested on less molecules than others, and downweighing extensively tested receptors. For a given molecule (*m*) receptor (*r*) interaction producing binary binding outcome *y*, identified from screen *type*, the final sample weights are given by the product of the three terms:

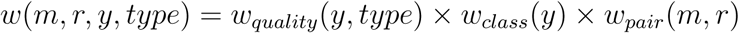

Where each of the terms represent the sample quality, label imbalance and pair imbalance weights.

#### Sample/Data quality weights

Due to the cost of conducting an experiment and ensuring high confidence on responsiveness, standard procedure for testing molecule-receptor pairs entails the following:

- First, a noisy *primary screening* is performed where many different molecule-receptor pairs are tested with one injection at a single concentration.
- Responsive pairs from the primary screening are tested at several concentrations in a *secondary screening*
- Pairs that are still considered responsive are testing through precise *tertiary screening (dose-response curves)* at multiple concentrations and multiple injections to form replicates.

As a result, considering the sample quality of a given molecule-receptor sample data point is vital to avoid fitting corrupted, noisy labels, affecting downstream generalization. Thus, prior work utilizes an estimated probability 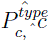 based on (*33*) where many pairs are tested across all three types of experiments to gain an empirical conditional probability of a given label being true/false (i.e: matching the dose-response label).

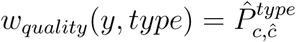

In this regime, dose-response screens (*EC*_50_) are denoted as the true label, with a conditional probability of 1. Please refer to (*9*) for the screening-specific screening conditional probabilities. **Class imbalance:** As most odorant-receptor pairs are non-responsive, the weighted loss uses an additional term which uses standard class weights based on the imbalance ratio to account for this. We also experiment with a softened version of the class imbalance term, where each descriptor’s contribution to the loss is weighted by a factor of *log*(1 + *w_class_*(*y*)). This is such that rarer descriptors are given a higher weighting without encouraging the model to over-predict compound-protein interactions as active. However, in our ablations we find that it does not have a significant difference in performance on the M2OR test set. Thus, we choose to utilize the model trained with the vanilla label imbalance ratio loss term to generate OR binding profiles for percept prediction.

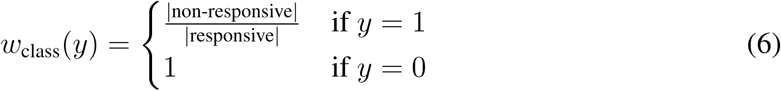

#### Pair imbalance

In addition to the class imbalance, there also exists a seperate type of data imbalance, where some olfactory receptors have been trained across many diverse molecules, while some have been tested across a few specific odorants. To avoid biasing across the uneven distribution of experiments, Hladis et. al. propose a weighing scheme to upweight less known receptors and down-weight extensively tested ones (*9*). Let |*M* (*r*)| be the number of molecules tested for a given receptor *r*, and |*OR*(*m*)| be the number of receptors tested for a given molecule *m*, where *K* is a constant (we follow the original work in setting *K* = 100).

The pair imbalance weight is then set to the logarithm of the harmonic mean:

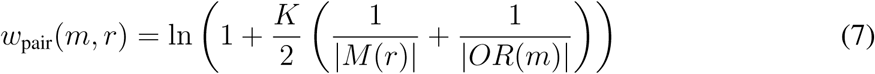

### Using olfactory receptor binding profiles for percept prediction

We investigate the use of the odorant-receptor model’s in-silico computed predictions as supplementary features for predicting the percepts of an odorant. Here given any arbitrary predictor of odorant-receptor and binding, for each molecule in the GS-LF dataset (5862 molecules), we generate predicted OR logits for the 1237 OR sequences in M2OR, sorted by of number of activated odorants (broadness). We repeat the same procedure for the 845 OR sequences in HORDE (despite MolOR not having trained on these sequences). For the final GS-LF benchmark of infilling the molecular features with OR logits (binarized through sigmoid activation and round operation), we train an odorant-receptor model (herewise dubbed “MolOR”) using the cross-attention style architecture described above on the M2OR odorant-receptor set. We modify the random splits on M2OR to be 90/10 to maximize training data but preserve some validation data for early stopping, in contrast to the original 80/10/10 train/val/test splits used to measure test performance on M2OR earlier.

The only modification to the original percept model architecture using vanilla GCNs for multi-task percept prediction is the concatenation of a binary vector of size (`M2OR # OR sequences = 400, HORDE # OR sequences = 845`). Here, any element *activation_i,j_* in the odorant-receptor matrix used is a binarized prediction of whether the odorant molecule *m_i_* in GS LF is activated by receptor *r_j_*, for *i* ∈ (1, 5862)*, j* ∈ `num OR sequences`. This binary vector is concatenated to the post-aggregated molecule vector, the output of the Weighted-Sum-and-Max aggregation performed on the node features. The modified molecule vector with the OR binding profile is passed to the same MLP prediction head to predict a binding score. The number of OR sequences to compute OR predictions for and concatenate to the molecular vector is treated as a hyperparameter, where we measure the information gain of scaling over how many OR sequence binding predictions we provide as context, going from more broad ORs (more activation data) to highly specific variant OR sequences. We ablate over the following values: `number receptor sequences` ∈ {0, 5, 10, 20, 40, 100, 400, max OR sequences}, but use the original trained MolOR model with 80/10/10 train/val/test random splits. Results for this experiment can be found in **Table S1**.

#### Training details

We employ the same hyperparameters as expanded on in the vanilla GCN for percept prediction, but modify the MLP prediction head to take in an input size of *gnn hidden feats*+*OR activations*. All other training details remain the same as the baseline architecture.

## Supplementary Information for

### Ablation Studies

**Table S1:** Effects of integrating more OR activation predictions from MolOR on GS-LF percept prediction.

| # OR Sequences | M2OR (Weighted) | HORDE (Weighted) | Stat. Test against Baseline |
| --- | --- | --- | --- |
| 0 (Baseline) | 87.04 ± 0.48 | 87.04 ± 0.48 | — |
| 5 ORs | 86.80 ± 0.44 | 87.29 ± 0.39 | $t = 2.84$, $q = 0.066$ |
| 10 ORs | 86.91 ± 0.28 | 87.19 ± 0.21 | $t = 1.96$, $q = 0.141$ |
| 20 ORs | 87.03 ± 0.76 | 87.33 ± 0.30 | $t = 4.67$, $q = 0.027^*$ |
| 50 ORs | 87.24 ± 0.17 | 87.27 ± 0.24 | $t = 3.48$, $q = 0.044^*$ |
| 100 ORs | 86.91 ± 0.44 | 87.17 ± 0.38 | $t = 0.63$, $q = 0.560$ |
| 400 ORs | **87.57 ± 0.49** | 87.24 ± 0.31 | $t = 5.05$, $q = 0.027^*$ |
| All ORs | 87.35 ± 0.36 | **87.62 ± 0.33** | $t = 4.43$, $q = 0.027^*$ |

Statistics are paired t-tests vs baseline ( $n_{\text{OR}}=0$ ), corrected for multiple comparisons using Benjamini-Hochberg FDR across 7 ablations. \* $q < 0.05$ . We additionally report a Jonckheere-Terpstra test for monotonic trend across all ablation OR levels ( $p = 0.010$ ).

We measure the contribution of using progressively more synthetic predictions of odorant-receptor binding for the percept prediction task. In Table S1. we show, using the sample-weighted MolOR model, that concatenating a binary vector of M2OR/HORDE receptor activation’s from the MolOR model to the aggregated molecular embedding from the GNN results in additional predictive power as the # of OR predictions scales to the full sets of receptors. We ablate the number of OR predictions given per sample, starting with only the top 5 OR sequences, to all 845-1237 sequences. We find that using the HORDE sequences (only human ORs rather than mammalian) offers more consistent improvement’s in performance, although both aid in percept mapping. For M2OR, the sequences are sorted in terms of broadness, or amount of positive active odorants (*14*). We find that the percept prediction performance steadily scales, and utilizing the top 845 OR logits’ synthetic predictions in HORDE produces a 0.6% increase to AUROC across 5 seeds.

We also measure the contribution of both a) the cross attention module over *per-residue embeddings* and b) the contribution of evolutionary representations learned by protein language models. Utilization of per-residue embeddings, indicated by the MolOR models (both naive, weighted), results in a over 10% improvement on AUC, when compared to the baseline, concatenating the equivalent mean-aggregated ESM embedding. This demonstrates that the cross-attention step to fuse information between both node-level odorant features, and residue-level receptor features, enables the model to learn more informative features in aggregating the residue features into a fixed size vector. For b), we show that using a pre-trained ESM model to generate the per-residue OR embeddings for MolOR results in a 3.8% boost over using a randomly initialized ESM model to infer residue-level features for the receptor representation. This demonstrates that the evolutionary representations captured by large-scale pLMs benefits the MolOR architecture in successfully matching odorants to their receptors.

### Receptor activation specificity

To address whether the observed percept-receptor associations exceed random expectation, we constructed a null distribution by randomly permuting percept labels across molecules while preserving overall label frequency. For each of ten percepts (five narrow: *meaty, sulfurous, nutty, roasted, smoky*; five broad: *fruity, floral, sweet, green, fresh*), we counted the number of receptors showing statistically significant association (Fisher’s exact test, Bonferroni-corrected p *<* 0.05) in both the observed and permuted datasets, repeating the permutation 100 times.

All five narrow percepts showed significant receptor associations that far exceeded the null expectation (all p *<* 0.01): nutty (28 significant receptors), roasted (24), meaty (22), smoky (12), and sulfurous (7), compared to a null mean of 0 in each case. In contrast, three of five broad percepts (fruity, green, fresh) showed zero significant receptor associations—consistent with the null distribution—suggesting these general odor categories arise from distributed receptor activation patterns rather than specific receptor subsets. We present these results in Figure S4, which directly compares observed versus null receptor associations for each percept.

**Figure S1:**
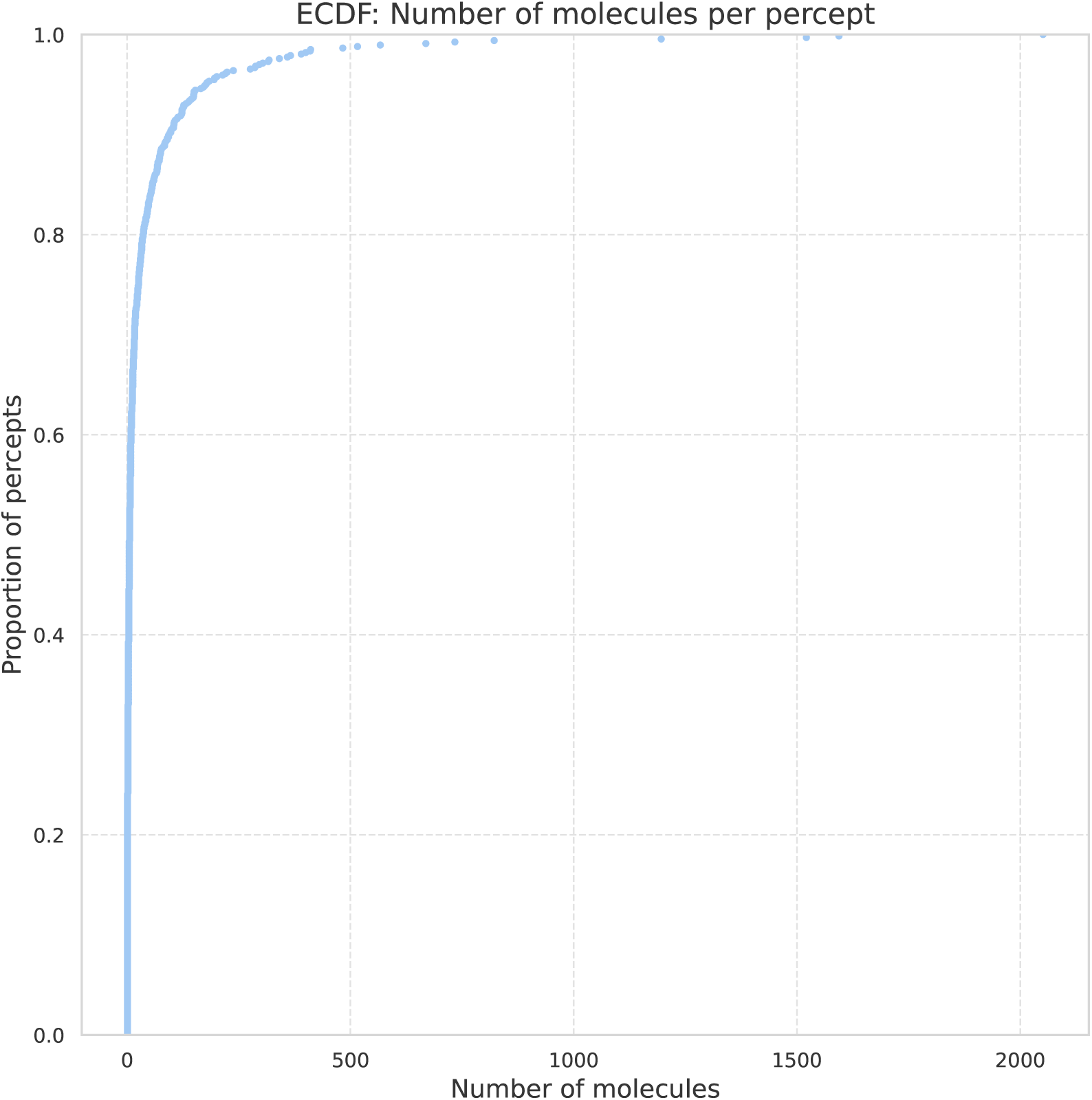
Goodscents-Leffingwell percept ECDF, demonstrating the imbalanced nature of the dataset. A few select percepts (sweet, fruity, vanilla) dominate majority of the labelled data, outlined by the few percepts with *>* 500 molecules).

**Figure S2:**
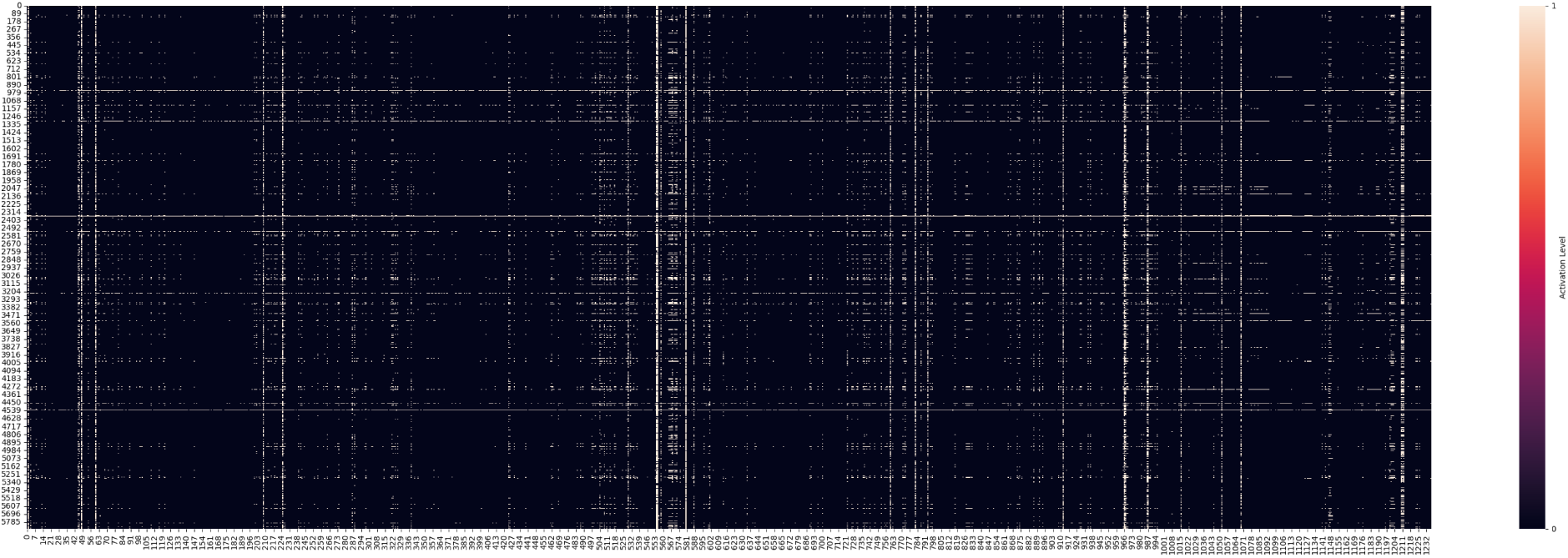
Plot of all predicted OR activations across the M2OR database, against GS-LF odorants (5862 odorants × 1237 OR sequences. Activations are predicted using the unweighted model.

**Figure S3:**
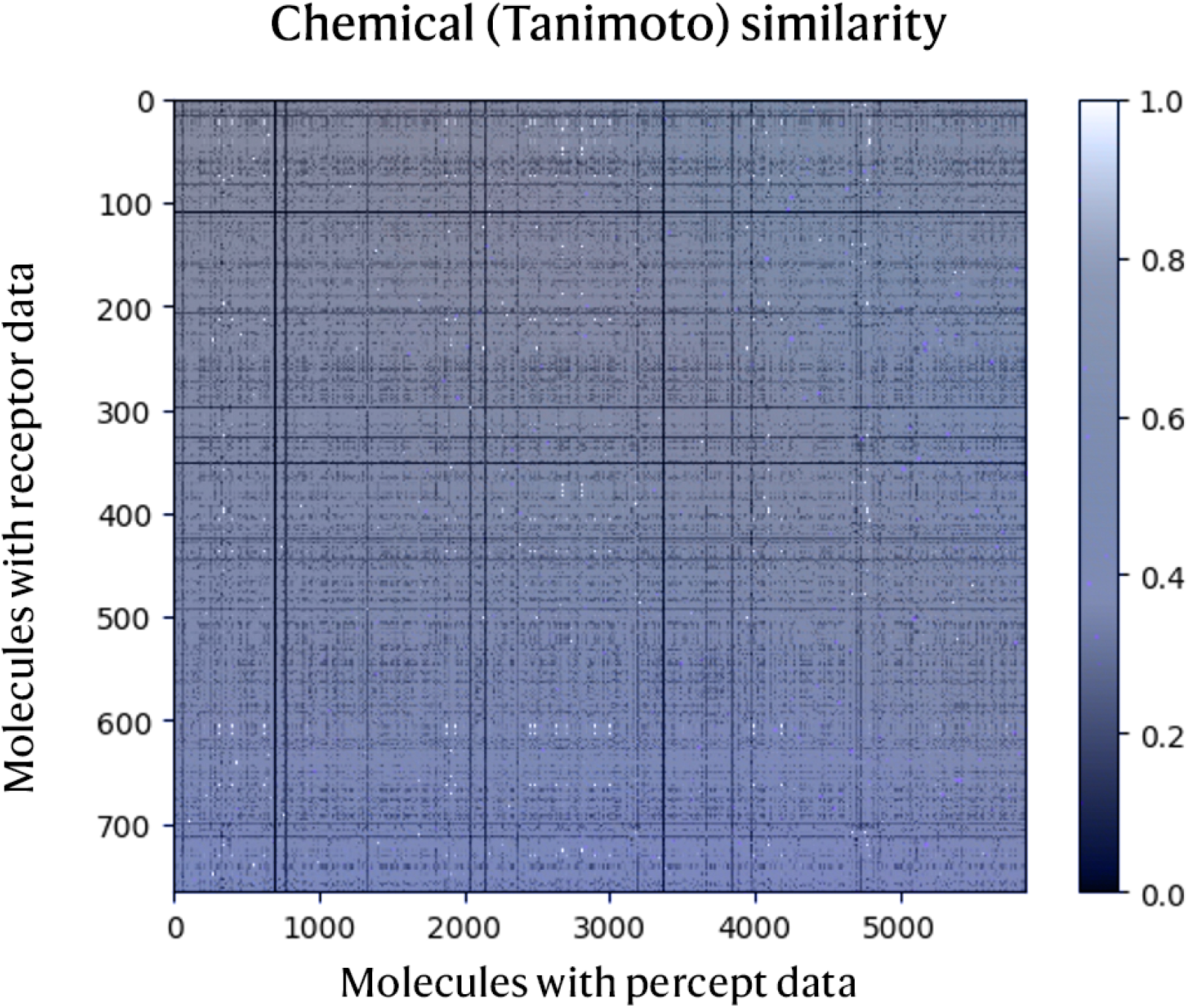
We see that there exists scarce end-to-end matches for mapping odorant to receptor interaction to perception, based on pairwise Tanimoto similarity scores in both M2OR and our assembled GS-LF dataset. There are no molecules that are in both datasets.

**Figure S4:**
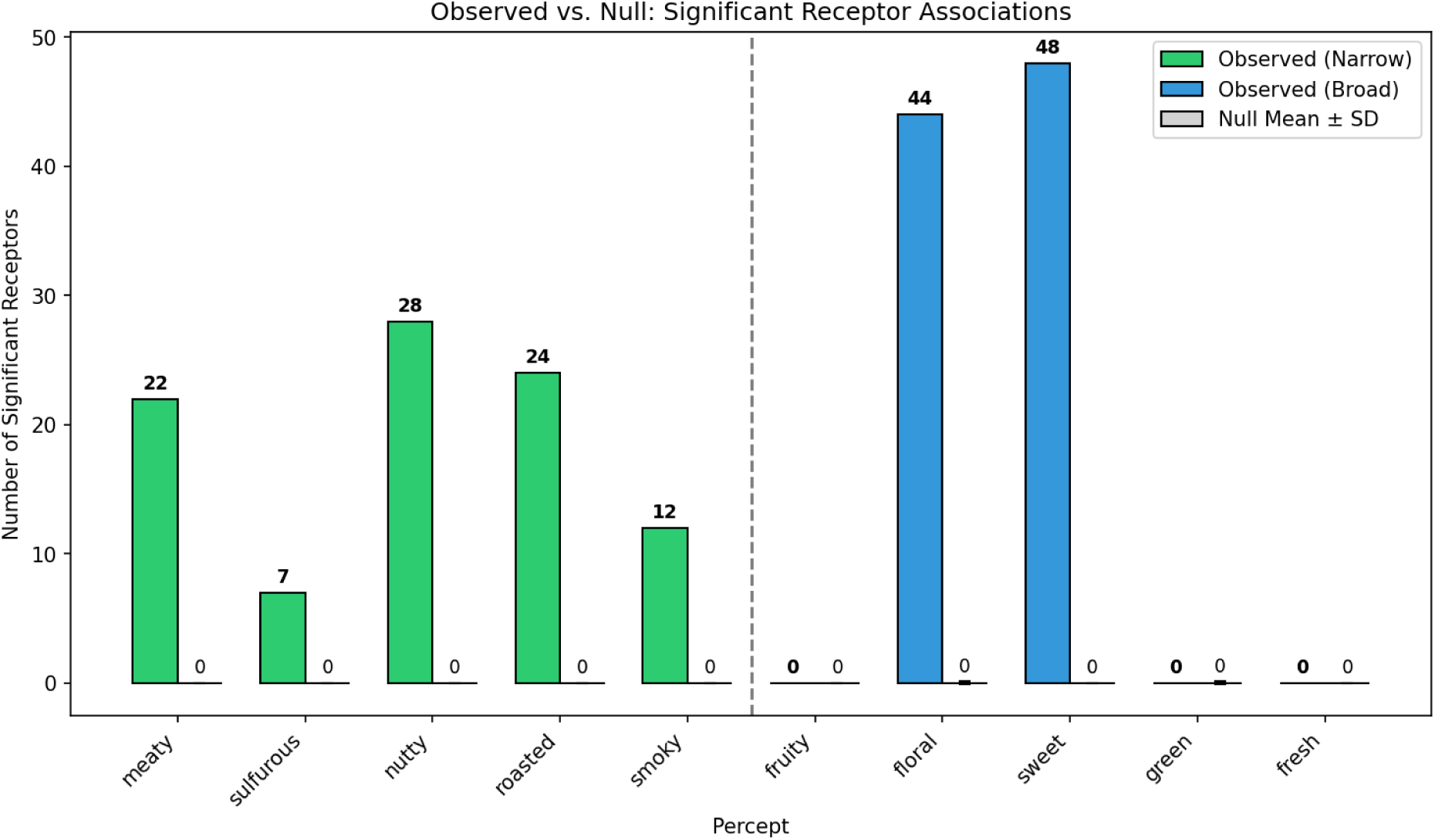
Percept-receptor associations exceed random expectation for narrow but not broad percepts. Here, a paired bar chart comparing the number of significantly associated receptors (Fisher’s exact test, Bonferroni-corrected p ¡ 0.05) between observed data (colored bars) and a null distribution (right bar for each individual percept) generated by shuffling percept labels across molecules (gray bars, mean ± SD from 100 permutations). Narrow percepts (green; meaty, sulfurous, nutty, roasted, smoky) show 7–28 significant receptor associations compared to 0 expected by chance, while three out of the five broad percepts (fruity, green, fresh) show no significant associations (fruity, green, fresh). Numbers above bars indicate the count of significant receptors.

## Notes

### Summary of Updates

Clarifications and ablations in response to reviewers

